# Predicting nanocarrier permeation across the human intestine *in vitro*: Model matters

**DOI:** 10.1101/2024.03.08.584089

**Authors:** Nathalie Jung, Jonas Schreiner, Florentin Baur, Sarah Vogel-Kindgen, Maike Windbergs

## Abstract

For clinical translation of oral nanocarriers, simulation of the complex intestinal microenvironment is crucial to evaluate interactions and transport across the intestinal mucosa for predicting the drug’s bioavailability. However, permeation studies are often conducted using simplistic cell culture models, overlooking key physiological factors such as tissue composition, morphology, and additional diffusion barriers as constituted by mucus. This oversight may potentially lead to an incomplete evaluation of the nanocarrier-tissue interactions and an overestimation of permeation. In this study, we systematically investigated different 3D tissue models of the human intestine under static cultivation and dynamic flow conditions with respect to tissue morphology, mucus production, and their impact on nanocarrier permeation. Our results revealed that the cell ratio between the different cell types (enterocytes and goblet cells), as well as the choice of culture conditions, had a notable impact on tissue layer thickness, mucus secretion, and barrier impairment, all of which were increased under dynamic flow conditions. Permeation studies with polymeric nanocarriers (PLGA and PEG-PLGA) elucidated that the amount of mucus present in the respective model was the limiting factor for the permeation of PLGA nanocarriers, while tissue topography represented the key factor influencing PEG-PLGA nanocarrier permeation. Furthermore, both nanocarriers exhibited diametrically opposite permeation kinetics in a direct comparison to soluble compounds. In summary, these findings reveal the critical role of the implemented test systems on permeation assessment and emphasize that, in the context of preclinical nanocarrier testing, the choice of *in vitro* model matters.

## Introduction

Many nanocarrier formulations designed for oral administration fail to accomplish the transition from preclinical development to clinical application due to their inability to effectively navigate the complex microenvironment of the intestinal mucosa to reach their target site. Rather than a flat layer of epithelial cells, the mucosa constitutes an intricate three-dimensional tissue comprising a variety of specialised cells and lined with a mucus layer, posing an additional diffusion barrier for nanocarriers. The therapeutic efficacy of oral formulations hinges on their ability to traverse the mucosal barrier by evading entrapment within the mucus layer and effectively crossing the underlying cell barrier. During the rational design of nanoparticulate formulations, it is therefore crucial to perform functional characterisation assays by investigating the interaction between the nanocarriers and biological tissue. Following technical advancements and the broad acceptance by regulatory agencies, the preclinical evaluation of novel therapeutics and their interaction with human cells and tissues is increasingly performed in *in vitro* systems rather than animal models^16–19^. The *in vitro* approach offers a controlled environment for the characterisation of pharmacokinetic and pharmacodynamic processes and allows, in combination with *in silico* tools, for the prediction of the *in vivo* performance of the formulations^20,21^. Cell-based assays are implemented to assess the biocompatibility of the formulation by measuring metabolic activity or cytotoxic effects and are further used to observe the biological function of the material and encapsulated drug^22–24^. In the context of oral formulations, the efficiency of targeting approaches or the uptake into cells might be investigated, but in general, the ability of the nanocarrier to cross the biological barrier of the intestinal mucosa is used to estimate their oral bioavailability as a measure for their performance^25–28^. The desire to accelerate nanocarrier formulation optimisation prompts many researchers to combine formulation development and biological testing in the same laboratory to allow for rapid adaptions. To achieve this, simplified cell monolayer permeability studies are often employed, particularly Caco-2 assays recommended by the FDA in an ICH harmonised guideline for biopharmaceutics classification system-based biowaivers^29^. The standardized cultivation of Caco-2 monocultures has made cell culture studies more accessible to researchers in fields such as chemistry and pharmaceutical technology who may not have extensive experience with handling biological specimens. Despite their usefulness, these simplified models neglect key physiological factors of the intestinal epithelium that could substantially impact on the interaction with nanocarriers. The exclusion of the mucus barrier, villi topography, and fluidic forces can distort fundamental assumptions about nanocarrier-tissue interactions, leading to inaccurate assessments of drug bioavailability. Strategies to overcome these limitations comprise the integration of mucus producing cell types, induced pluripotent stem cells or patient-derived material, and their cultivation in microfluidic chips, resulting in the formation of complex three-dimensional micro-tissues^30–35^. While these approaches offer the potential to create test systems with improved physiological relevance, they require a high level of expertise in the fields of cell biology as well as engineering and are currently limited by high production costs and low throughput. Efforts to strike a balance between manageable *in vitro* models and models with a higher degree of physiological relevance have led to the development of co-cultures of the enterocyte cell line Caco-2 and the mucus-producing cell line HT29-MTX, cultured on permeable membrane inserts^36,37^. Such test systems have gained widespread attention in nanocarrier research due to their relative ease of use and ability to provide insights into the role of mucus in intestinal barrier function. Despite their popularity, there is significant variability in the used cell ratios and culture conditions across different laboratories. Besides monocultures of both cell lines, the ratios of 90:10, 80:20, 70:30, 75:25, and 50:50 (Caco-2:HT29-MTX) are frequently reported in literature^38–42^. As a result, the impact of these variations on the properties of the resulting tissue, particularly the barrier integrity, secreted mucus amount, and the implications on nanocarrier interaction, remain poorly understood. While there is a range of reference literature on permeation properties of model drugs, small molecules, and fluorescent dyes across different *in vitro* models, our understanding of the interaction and permeation of nanocarriers across tissues remains limited^43–46^. To address the current knowledge gap concerning the permeation of nanocarriers across the intestinal epithelium *in vitro*, we conducted a systematic evaluation of commonly used human cell line-based tissue models by examining four different ratios of Caco-2 and HT29-MTX cells under both static cultivation and dynamic flow conditions. With the aim to implement a dynamic cultivation that can be realised independently from microfluidic setups, we opted for an attainable approach based on orbital agitation. We characterised the resulting tissue models in terms of tissue topography, barrier properties, and mucus secretion to obtain a comprehensive understanding of physiological changes. The consequential impact on the permeation of polymeric nanocarriers was subsequently assessed using two well-established nanocarrier formulations based on poly(lactic-co-glycolic acid) and polyethylene glycol.

## Results and Discussion

The present study provides a comprehensive evaluation of commonly utilized human cell culture models for assessing the permeation of nanocarriers across the human intestinal epithelium. While these models are well-established for evaluating the permeability of solute drug molecules^39,43,45–47^, the influence of cell composition, tissue topography, and mucus secretion on nanocarrier permeation remains poorly understood. Therefore, we have adopted a systematic approach to thoroughly characterise eight different *in vitro* models with respect to their physiological properties and the resulting implications on nanocarrier permeation. The human cell lines Caco-2 (enterocyte-like) and HT29-MTX (goblet cell-like) were seeded in different ratios (100:0, 90:10, 75:25, 0:100) to observe effects in monocultures as well as co-cultures with physiological ratios found in the human small intestine (90:10) and colon (75:25). The tissue cultures were cultivated in Transwell^®^ inserts for 21 days under static or dynamic conditions, by placing the culture plates on the incubator shelf or an orbital shaker, respectively (**Figure 1**).

**Figure 1.**
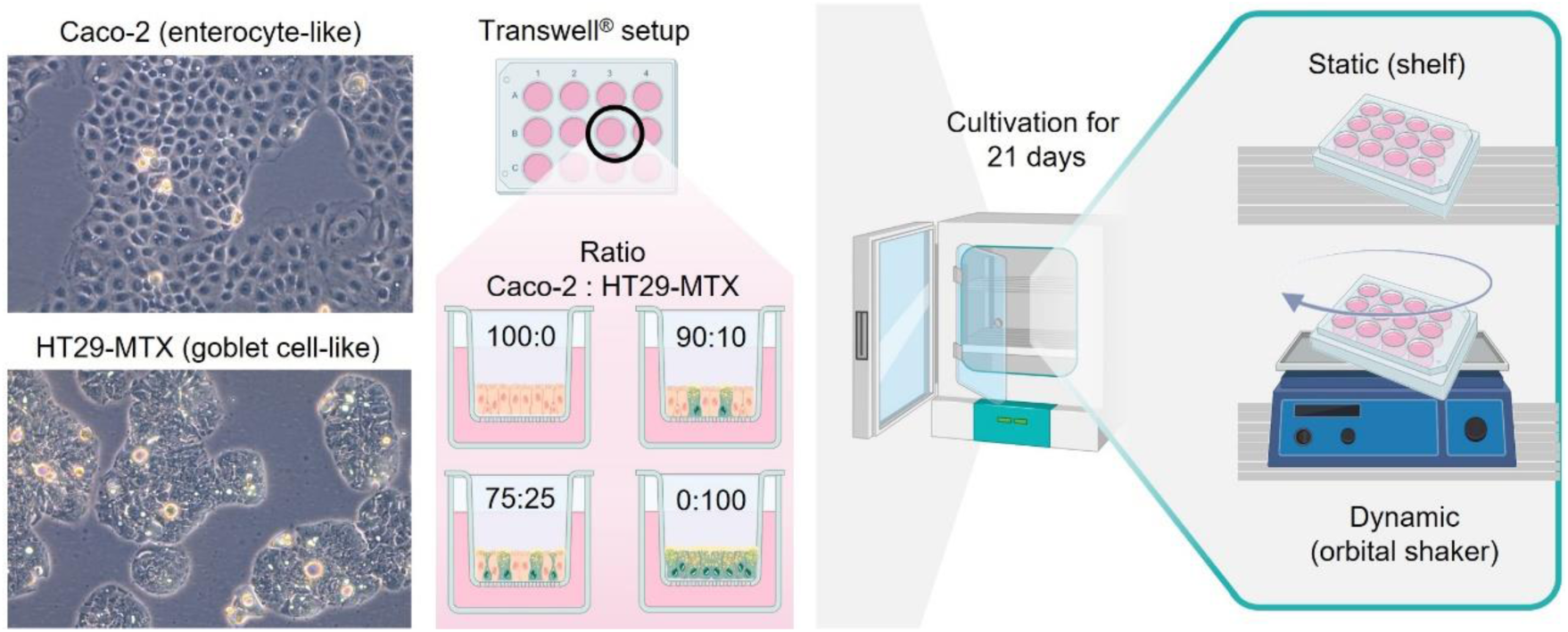
Schematic overview of the cultivation procedure of intestinal *in vitro* models. Enterocyte-and goblet cell-like cell lines (magnification 20x) were seeded in four different ratios and cultivated for 21 days under static (incubator shelf) or dynamic (orbital shaker) conditions.

### Tissue morphology

*In vitro* models of the human intestinal epithelium display an even, planar surface morphology when cultured under static cultivation conditions and undergo a remodelling of the tissue topography when exposed to shear forces under dynamic cultivation conditions^48–50^. The subsequent changes in tissue layer thickness, three-dimensionality, and surface area will consequentially impact the diffusion of drug molecules and nanocarriers^51^. To evaluate the impact of static and dynamic culture conditions on the tissue topography of the *in vitro* models used in the present study, we conducted histological analysis by alcian blue/periodic acid–Schiff (PAS) staining (Figure 2A). The staining method enabled the discrimination between nuclei (dark purple), cytosol (light purple), and acidic mucins (bright blue). Mucus-producing HT29-MTX cells were distinguished from Caco-2 via the presence of acidic mucins^52,53^. The number of mucin-positive cells correlated with the initial seeding ratios of Caco-2 and HT29-MTX cells, showing no mucins in Caco-2 monocultures and increasing amounts of mucin-positive cells in the 90:10, 75:25, and HT29-MTX monocultures. Although a notable layer of mucus was visible on top of HT29-MTX-containing *in vitro* models during the culture period, acidic mucins were solely observed within goblet cells in histological staining. Secreted mucins were most likely removed during the multiple washing steps involved in the histological sample preparation and staining procedure. The ratio of the two cell types further influenced the thickness of the cell layer in the respective cultures under static conditions (Figure 2B). Enterocytes exhibited a cuboidal morphology, while goblet cells showed a more columnar shape with mean epithelial layer thicknesses of 11.19±5.14 and 25.22±5.67 µm, respectively (larger magnifications see Figure SI1). The co-cultures of both cell types presented an intermediate epithelial layer thickness (13.62±7.00 µm for 90:10, 16.07±6.53 µm for 75:25) between the respective monocultures. The shear stress produced by the circular motion during dynamic cultivation induced a notable change in tissue morphology, leading to a more columnar cell morphology in both cell types. A significant increase in tissue layer thickness was observed for all cultures compared to statically cultivated models, measuring mean thicknesses of 47.85±19.71 µm (100:0), 46.04±23.88 µm (90:10), 59,96±27.54 µm (75:25), and 90.69±23.71 µm (0:100). Further, the applied shear stress promoted the formation of three-dimensional protrusions from the epithelium, which ranged from 80-120 µm in height and approximately 100 µm in width, independent of the cellular composition of the respective model. In cultures containing Caco-2 cells these protrusions comprised multiple cell layers and resembled the shape of intestinal villi. In HT29-MTX monocultures, the protrusions formed interconnected structures above the basal epithelium, resembling a continuous surface layer. The observed protrusions reached approximately 10 to 25% of the length of villi of the human small intestine, which typically measure 400 to 1000 µm^54^. However, the diameter of the protrusions observed in the *in vitro* models was similar to the dimensions of villi of the human small intestine^55^. Compared to microfluidic chip-based tissue models of the small intestine, the length, width, and density of the reported protrusions seem to correspond to those of the dynamically cultured *in vitro* models analysed in this study, although exact measurements are lacking in the cited literature^30,48,56,57^.

**Figure 2.**
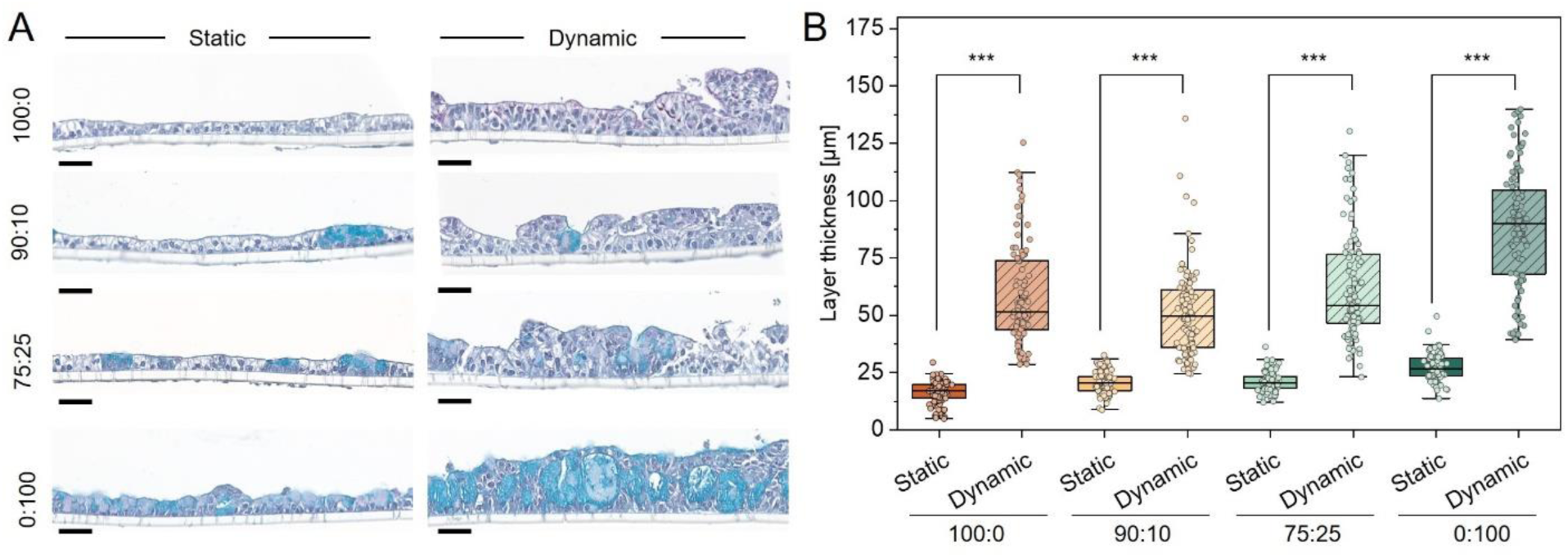
Alcian blue/PAS staining of histological sections of *in vitro* tissue models cultured under static or dynamic conditions (A). Layer thickness of epithelial tissues (B). Scale bars: 50 µm.

To develop a better understanding of the tissue architecture, we performed scanning electron microscopy allowing the visualisation of the epithelial surface of the *in vitro* tissues (**Figure 3A**). In statically cultured *in vitro* models, the surface showed a flat topography without any notable protrusions, confirming the findings from histological evaluation. In contrast, scanning electron micrographs of the dynamic cultures displayed an increasingly structured surface from Caco-2 monocultures to 75:25 cultures. Based on electron scanning micrographs, it becomes apparent that the number of protrusions increased notably in *in vitro* models with higher proportions of HT29-MTX cells. Profilometry based on white light reflectance further highlights the topographic changes between static and dynamic cultivation (**Figure 3B**) and allows for better visualisation of the height differences in goblet cell monocultures, which exhibited a planar epithelial surface morphology under both culture conditions. While the observed protrusions in dynamic co-cultures seemingly took the shape of flattened villi in histological cross-sections, the acquired micrographs revealed an elongated, stretched, and interconnected morphology in many of the structures. In literature, the analysis of tissue topography of microfluidic chip-based models mainly includes cross sections in form of histological or (immuno)fluorescence staining, depicting villi-like structures^57,58^. Only occasionally, publications include light microscopic images or electron micrographs that provide more thorough structural investigations of the surface morphology, showing similarly shaped protrusions^48^. We can assume that the orbital shaking induces comparable surface morphogenesis compared to microfluidic chip-based models and therefore represents a more attainable approach for modelling the three-dimensional surface of the intestine *in vitro*. Although the villi-like structures do not match the exact dimensions of the *in vivo* counterparts, using three-dimensional intestinal models might greatly influence the permeation of soluble drug molecules as well as nanocarriers due to the change of available surface area.

**Figure 3.**
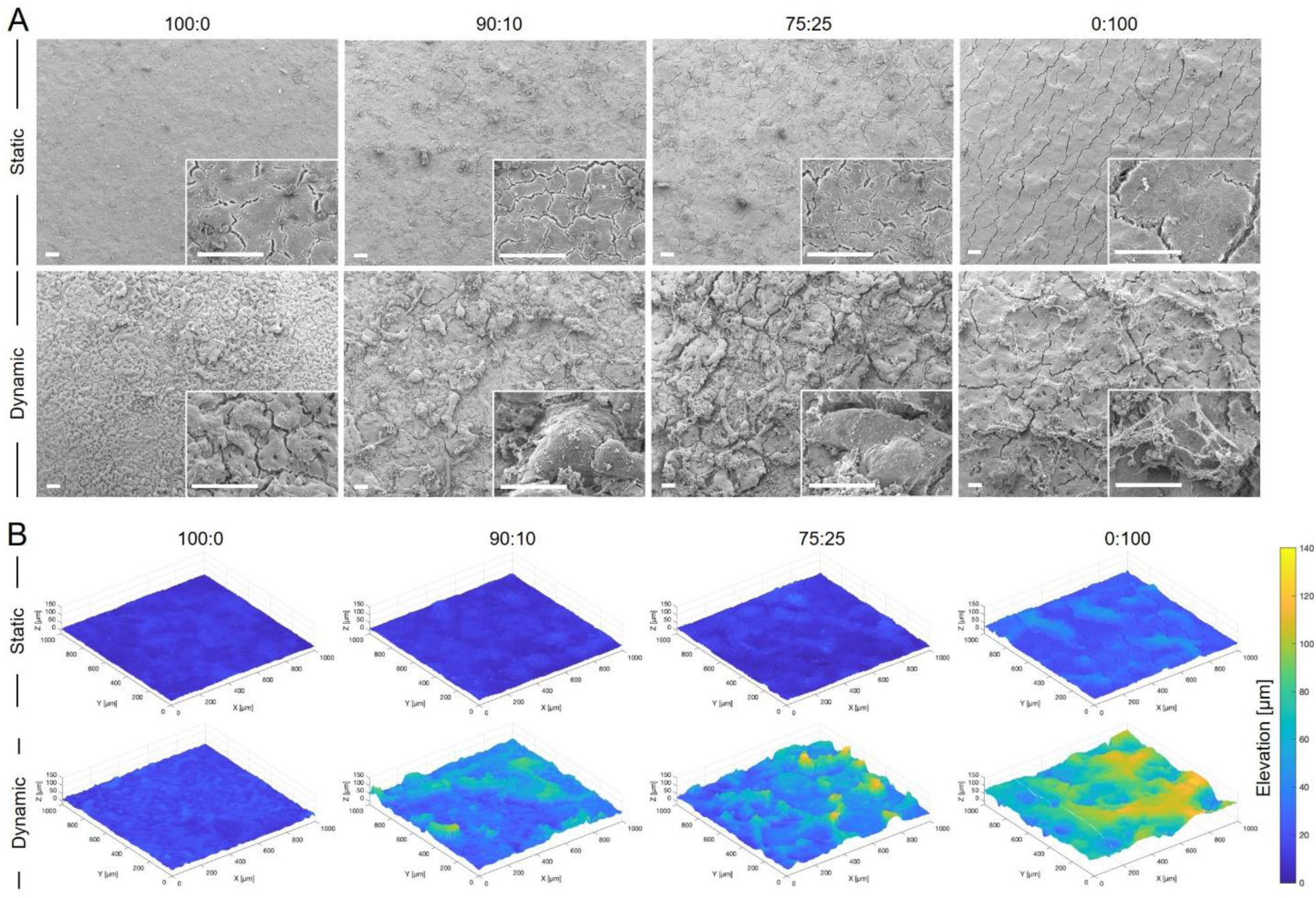
Scanning electron micrographs (A) and topography maps (B) of the surface of statically and dynamically cultivated *in vitro* models. Scale bars: 100 µm.

### Mucus secretion

The human intestinal cells are protected by a mucus lining that varies in thickness and poses an additional permeation barrier for orally administered drugs as well as nanocarriers. To account for this additional barrier, goblet cells (HT29-MTX) were integrated in *in vitro* models in differing amounts. We chose ratios of 90:10 and 75:25 (Caco-2:HT29-MTX) to mimic the cellular composition of enterocytes to goblet cells in the human small intestine and the colon epithelium, respectively^59,60^. To evaluate the impact of Caco-2 to HT29-MTX ratio and the influence of dynamic cultivation on mucus secretion, we quantified the amount of acidic mucins via *in situ* alcian blue staining. By varying the ratio of enterocytes to goblet cells in the models, an increase in mucus was observed via the presence of dark blue spots within the cell layer (Figure 4A). Upon dynamic cultivation, these mucin-rich spots grow in size and abundance in all cultures containing HT29-MTX cells. Although a notable layer of mucus on top of the epithelial layer was observed during cultivation, the secreted mucus layer was not visible in *in situ* staining. We suspect, that due to the multiple fixation and washing steps, secreted mucins were removed, and consequentially only intracellular mucins were stained and quantified. Absorbance-based quantification revealed an increase of 21.4% and 45.9% in mucus production in statically cultured 90:10 and 75:25 *in vitro* models, respectively, compared to Caco-2 monocultures (100:0) (**Figure 4B**). Mucus production in HT29-MTX monocultures (0:100) was increased by 72.9%. We found that the percentage of goblet cells in the statically cultured *in vitro* models (10%, 25%, 100%) was not matched by an equal increase in produced mucus. The dynamic cultivation of tissue models had a small effect on mucin production in 100:0 (+14.4%) and 90:10 cultures (+16.8%) but showed a more pronounced increase in models containing higher percentages of goblet cells. Dynamic cultivation resulted in an average increase in mucin production of 55.5% in 75:25 and 70.6% in 0:100 cultures compared to their static counterparts, respectively. Filamentous remainder of this mucins can also be observed in the scanning electron micrographs (Figure 3A). Overall, the increase of mucin secretion was notably more pronounced under dynamic conditions and is consistent with previous studies performed in chip-based *in vitro* systems^49^. However, with respect to the extent of the intestinal mucus layer *in vivo*, which can range from 20-100 µm in the small intestine and several hundred µm in the colon^61,62^, a comparison to *in vitro* models is difficult. Due to the quantification procedure, which involved fixation and washing steps, large amounts of mucus might have been removed from the epithelial surface, and the method did not allow for the determination of mucus layer thickness. Nevertheless, the grave differences in mucus secretion between static and dynamic culture underscore the need for critical model selection to characterise permeation abilities of nanocarriers *in vitro*.

**Figure 4.**
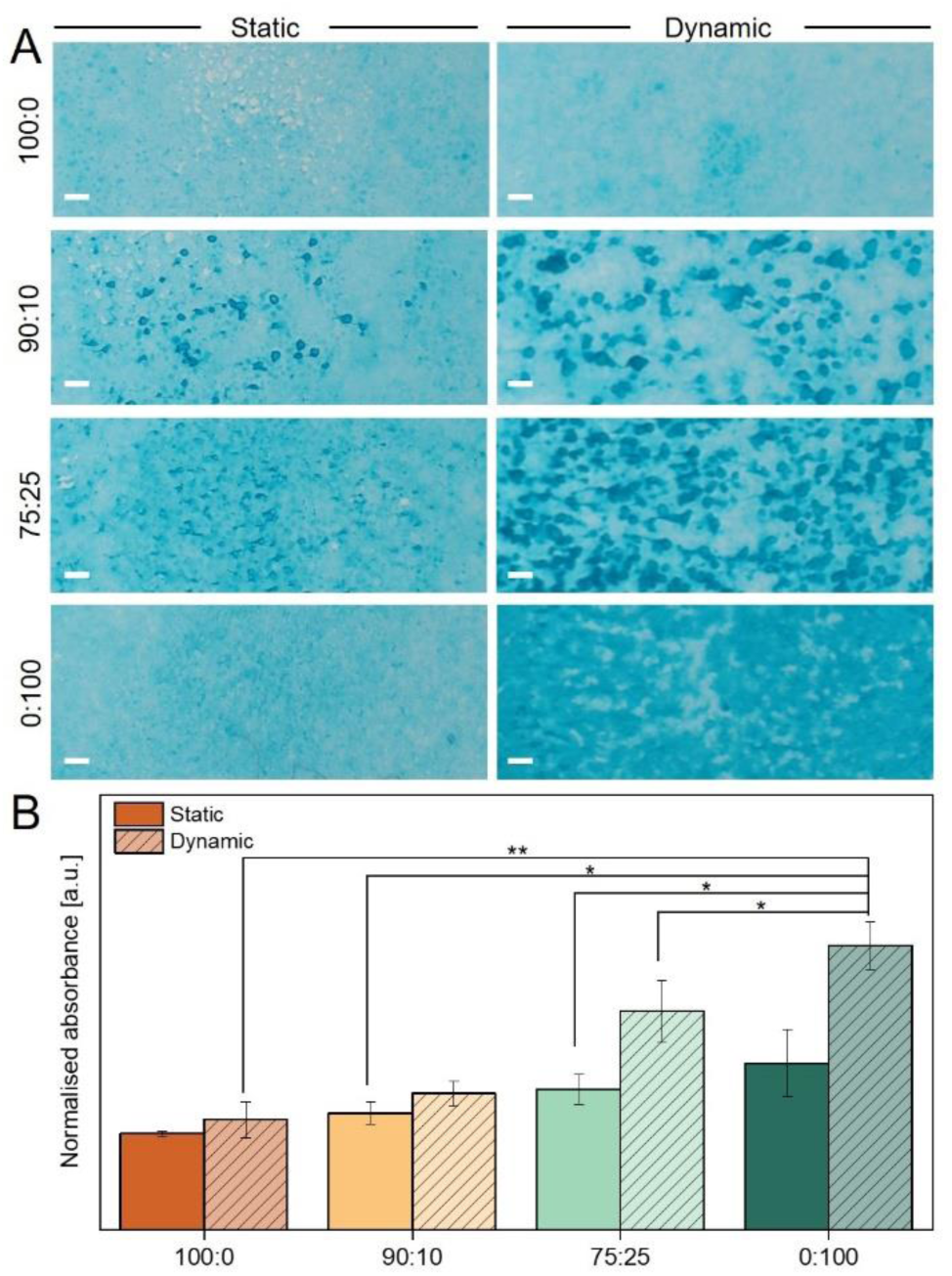
*In situ* alcian blue staining of acidic mucins (A) and semi-quantitative evaluation of mucus production (B). Scale bars: 100 µm.

### Epithelial barrier integrity

Below the mucus layer, the epithelium poses the next permeation barrier for nanocarrier formulations. For Caco-2-based cell culture models, a cultivation time of 21 days is required to allow the cells to differentiate into enterocytes, which exhibit microvilli on the apical surface and express tight junction protein complexes that are essential for barrier formation^63–65^. The presence of microvilli as an indicator for enterocyte differentiation was confirmed by scanning electron microscopy in all cell culture ratios and static and dynamic cultivation conditions (Figure 3A). To validate the formation of an intact epithelial barrier, the transepithelial electrical resistance (TEER) of the cell culture models was monitored over the course of the 21-day cultivation period (**Figure 5A-B**). Under static conditions, 100:0 cultures experienced an initial increase in TEER during the first 7 days, eventually reaching a plateau with a final TEER of 172±30 Ωcm^2^. The 90:10 and 75:25 co-cultures behaved similarly and exhibited comparable final TEER values (173±22 and 183±21, respectively) to the Caco-2 monoculture. In contrast, 0:100 cultures showed a slower but continuous increase in barrier integrity throughout the cultivation period and reached slightly higher TEER values compared to other static cultures. When subjected to dynamic conditions, TEER values of 100:0 cultures reached a maximum after just 7 days at 290±66 Ωcm^2^, followed by a decrease to 175±38 Ωcm^2^. 0:100 cultures again exhibited a slow and steady increase of barrier integrity under dynamic conditions but only reached a maximum of 81±23 Ωcm^2^ after 21 days. This value represented approximately 30% of the TEER observed in static 0:100 cultures. 90:10 and 75:25 co-cultures demonstrated an intermediate development between the two monocultures and exhibited a final TEER of 127±24 and 135±24 Ωcm^2^, respectively. Upon closer inspection of individual replicate measurements (Figure SI1), a more uniform development of TEER values of Caco-2 cultures under static conditions was observed, while larger deviations were measured under dynamic conditions. In contrast, HT29-MTX cultures showed the opposite characteristics in time-dependent barrier formation with higher deviations after 21 days in static culture. Overall, dynamic cultures containing goblet cells exhibited lower TEER values than their static counterparts at the end of the cultivation period. However, it is worth noting that these values corresponded to measurements performed on *ex vivo* human intestinal tissue, which exhibited TEER values around 100 Ωcm^2^, and therefore can be considered a physiologically relevant barrier for studying the permeation of nanocarriers across the intestinal epithelium^66,67^.

**Figure 5.**
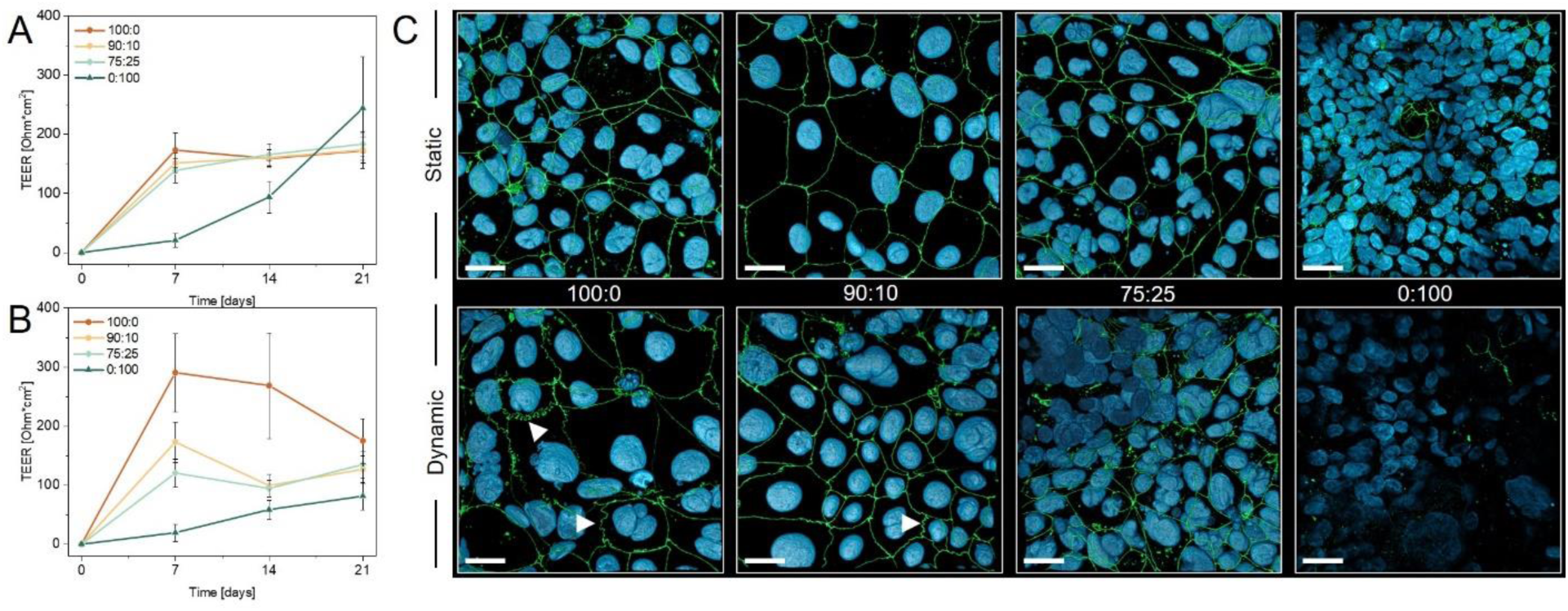
Transepithelial electrical resistance (TEER) of cell culture models under static (A) and dynamic (B) culture conditions. Fluorescence staining of the tight junction associated protein ZO-1 (green) and cell nuclei (blue) after 21 days in culture (C). Arrows indicate ribboning of tight junctions. Scale bars: 25 µm.

To further evaluate the integrity of the epithelial barrier in all *in vitro* models, we analysed the expression of the tight junction-associated protein zonula occludens 1 (ZO-1) as an additional marker (Figure 5C). While ZO-1 was clearly present in all investigated samples, statically cultured *in vitro* models exhibited a more homogeneous distribution of ZO-1 along the cell membrane. In dynamic models, ribboning and internalisation of ZO-1 was observed in multiple instances (indicated by arrows in Figure 5C), possibly contributing to the slightly lowered TEER values. The decrease in fluorescence intensity in co-cultures and 0:100 cultures, especially under dynamic conditions, was due to the increasing proportion of goblet cells, which inhibited efficient fluorescence staining due to higher amounts of secreted mucus.

### Tissue permeability

The permeability of the intestinal epithelial barrier of drugs of nanocarriers is dependent on the intricate interplay between both, the extent of the mucus layer and the properties of the cellular barrier with respect to tissue layer thickness and the development of cellular junctions^68,69^. After conducting a comprehensive characterisation of these factors in our intestinal *in vitro* models, we proceeded with transport studies to elucidate how tissue composition and culture conditions influence the epithelial permeability of model drug substances as well as of polymeric model nanocarrier formulations (**Figure 6**). First, we employed FITC-dextran-based leakage studies as an established procedure to determine the permeability of solute substances through tissue models. The mean apparent permeability coefficient (P_app_) for 4 kDa FITC-dextran is shown in Figure 6A. Under static conditions, FITC-dextran exhibited comparable P_app_ values in 100:0 monoculture and co-cultures (90:10 and 75:25), probably due to their similar physiological properties observed in earlier experiments (tissue layer thickness, mucus secretion, TEER). However, HT29-MTX monocultures showed higher P_app_ values of FITC-dextran despite increased layer thickness and TEER values. Similar discrepancies between TEER and P_app_ have been observed in different tissue models and highlight the inherent necessity of performing control experiments to assess the transport of soluble substances^70–72^. Under dynamic conditions, the P_app_ values increased notably for all cultures, although TEER values of 0:100, 90:10, and 75:25 cultures were comparable to their static counterparts. Dynamic 0:100 monoculture exhibited the highest increase. These data show that monocultures of HT29-MTX cells were more permeable than Caco-2 monocultures and co-cultures of both cell lines, which supports recent findings from a study performed by McCright et al., where similar cell ratios were compared after 21 days of static cultivation^43^. Further, we found that the dynamic cultivation of *in vitro* models overall induced higher permeability of the tissue barrier for solute substances. The impact of dynamic conditions on barrier permeability in intestinal *in vitro* models is not thoroughly characterised in current literature and yields conflicting results depending on the choice of cell source and cultivation setup. In a study from Schweinlin et al., primary human enterocytes were cultivated in a similar dynamic cultivation procedure with lower shear stress and no alteration in barrier permeability was observed^73^. In another study, where Transwell^®^-based Caco-2 cultures were compared to microfluidic chip-based systems, a higher permeability was measured using dynamic cultures, similar to the trends observed in the here presented study^74^.

**Figure 6.**
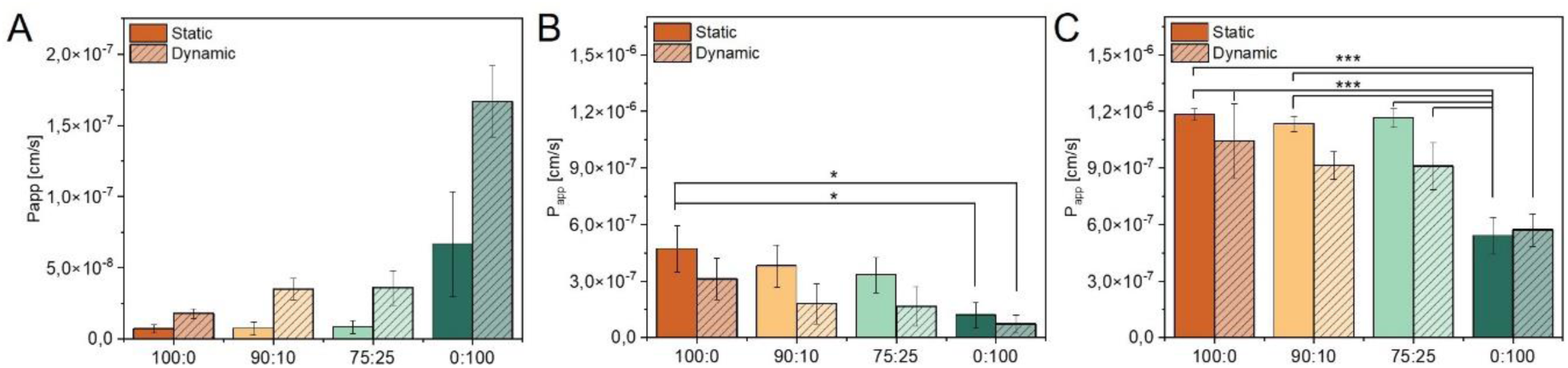
Apparent permeability (P_app_) of 4 kDa FITC-dextran (A), PLGA nanocarriers (B), and PEG-PLGA nanocarriers (C) determined via the transported mass across different *in vitro* models.

Given the complex barriers of the intestinal mucosa, which impact the permeability of substances based on their size and physicochemical properties, our study ultimately aimed to elucidate the influence of the choice of *in vitro* model on the permeation characteristics of nanocarriers. Therefore, we assessed the transport of nanocarriers across the different *in vitro* models using two types of commonly used polymeric nanocarriers in the field of drug delivery. We fabricated PLGA nanocarriers with an average size of 180.7±10.7 nm and a zeta potential of - 37.2±4.9 mV, as well as PEG-PLGA nanocarriers measuring 181.8±20.5 nm in size and having a surface charge of −36.4±4.2 mV (Figure SI3). These well-established model formulations were selected for the study based on the extensive knowledge and understanding of their characteristics and their potential for oral drug delivery^75–78^. The implementation of PLGA as more hydrophobic polymer and the addition of PEG as commonly employed hydrophilic surface modification allows for the evaluation of two nanocarrier formulations, each exhibiting distinct physicochemical surface properties^79,80^. We assessed the permeation kinetics of both formulations in the different *in vitro* models under static and dynamic conditions over 6 h. PLGA nanocarriers showed the highest P_app_ values in statically cultured 100:0 models, and a steady decrease of permeability was observed with increasing percentages of goblet cells in the respective *in vitro* models (Figure 6B). The consequential higher amount of produced mucus on top of the cell layer is known to pose an additional barrier for the PLGA nanocarriers^81^. Static 0:100 cultures showed a significant drop in PLGA nanocarrier permeation of approximately 75% decrease compared to static 100:0 cultures. The observed trends were even more pronounced when investigating the permeability of PLGA nanocarriers in dynamically cultured models. P_app_ values in dynamic *in vitro* cultures were decreased by 33%, 52%, 50%, and 39% in 100:0, 90:10, 75:25, and 0:100 compared to their static counterparts, respectively. A highly significant decrease in nanocarrier permeation could be observed between static 100:0 and dynamic 0:100 cultures. As we were able to show during the characterisation of the *in vitro* models, the dynamic cultivation not only increased the layer thickness of the respective model, but also promoted an increase in mucus secretion in all goblet cell-containing cultures. Particularly for polymeric nanocarriers without surface modifications, this increase in mucus presented a hard to overcome biological barrier and was most likely the main contributing factor in the reduced permeability^81,82^.

PEG-PLGA nanocarriers overall exhibited higher P_app_ values compared to PLGA carriers, due to the hydrophilic properties of PEG chains^11,83^. The permeation ability of nanocarriers in statically cultured 100:0, 90:10, and 75:25 models was comparable and showed only a small decrease in cultures with higher proportions of goblet cells. Dynamic cultivation of these models led to an average decrease of 14%, 20%, and 22% in comparison to their static counterparts, respectively. Since a lower P_app_ for PEG-PLGA nanocarriers was also observed in dynamic 100:0 cultures, which do not produce notable amounts of mucus, we suggest that the effect might be attributed to changes in the tissue morphology rather than the increase in mucus secretion during dynamic cultivation. This hypothesis was supported by confocal fluorescence images of the epithelial cell layer after the transport study (Figure SI4). In these images, nanocarriers were observed to accumulate within the inter-villus regions, resembling valley-like structures in the intestinal models. This accumulation suggests a potential reduction in the available surface area of the epithelium for active permeation processes. Moreover, 0:100 monocultures exhibited significantly lower nanocarrier permeation in static and dynamic models, indicating a permeation-reducing effect of mucus in *in vitro* models with percentages of goblet cells above 25%. Despite the increase in mucus secretion in 0:100 cultures under dynamic conditions, no notable differences in PEG-PLGA nanocarrier permeation between static and dynamic 0:100 cultures were observed. Interestingly, the layer thickness of these models did not appear to be an attributing factor to permeation abilities either, as dynamic 0:100 cultures would have been expected to experience a reduction in P_app_ values if this were the case.

Moreover, the differences in three-dimensional tissue architecture of dynamically cultured *in vitro* models might be attributed to differences between enterocyte-containing models, which exhibit a villi-like morphology, and 0:100 cultures, which showed a flat epithelial surface but an overall increased tissue layer thickness. However, the attribution of permeation inhibition to the available surface area in each model is difficult due to the lack of advanced imaging systems that allow for the real-time monitoring of permeation processes and requires further investigation.

To ensure the reliability of our findings regarding the permeation ability of FITC-dextran and nanocarriers, we conducted standard quality assessments during each experiment. This involved measuring the TEER values before and after the transport studies, as well as evaluating the cytotoxicity of nanocarrier formulations. These assessments were performed to validate that our observations were specific to the permeation ability of the substances and not influenced by toxicity-induced increased permeability of the *in vitro* models (Figure SI5).

### Conclusion

In the here presented study, we set out to elucidate how the selection of the *in vitro* tissue model impacts the assessment of nanocarrier permeation kinetics in oral drug delivery. In a systematic approach, we demonstrated that varying cell ratios (enterocytes to goblet cells) and cultivation conditions (static versus dynamic) have complex effects on key physiological properties of *in vitro* models of the human intestinal mucosa. The resulting variations in three-dimensionality, epithelial surface morphology, mucus secretion and barrier properties were found to significantly influence the permeation kinetics of nanocarrier formulations. Notably, depending on the carrier type different physiological features of the models were identified as individual rate-limiting factors. These findings show that the choice of *in vitro* model matters during the assessment of transepithelial permeation of nanocarriers, highlighting the critical need to carefully select *in vitro* models during formulation development. By raising awareness for the choice of a suitable model for predictive permeation testing of nanocarriers, this study might contribute to address a key challenge hindering the translation of nanomedicine formulations from preclinical development to clinical application.

## Materials and Methods

### Cell culture

Cultivation of intestinal *in vitro* models was performed using the Caco-2 clone C2BBe1 (LGC Standards, Teddington, UK) passages 50 to 64, and HT29-MTX-E12 (ECACC, Porton Down, UK) passages 52 to 66. The two cell lines were cultured separately in Dulbecco’s Modified Eagle Medium (DMEM) (Gibco, Carlsbad, US) supplemented with 10% foetal bovine serum (Sigma Aldrich, Darmstadt, Germany) and 1% non-essential amino acids (Gibco, Carlsbad, US) at 37°C and 5% CO_2_. Cells were subcultured once a week. *In vitro* models were seeded in Caco-2:HT29-MTX-E12 ratios of 100:0, 90:10, 75:25, and 0:100 with a seeding density of 5×10^4^ cells/cm^2^. Cells were seeded in 12-well Transwell^®^ inserts with a pore size of 3.0 µm (Falcon, Corning, US) and cultivated for 21 days under static (incubator shelf) or dynamic conditions. Dynamic cultivation was achieved by placing the cell culture plates on an orbital microtiter shaker (IKA, Staufen, Germany) at 175 rpm 24 h after seeding. The medium was replaced every 2 to 3 days.

### Histology

To assess the distribution of Caco-2 and HT29-MTX cells within the different *in vitro* models and visualise the cell layer organization, histological analyses were conducted on all cultures. *In vitro* models were washed twice with phosphate-buffered saline (PBS), pH 7.4 (===), and fixed by incubating the sample in a 3.7% buffered paraformaldehyde solution for 15 min at room temperature. Samples were dehydrated through an ethanol series of 70%, 80%, 95%, 100% ethanol, and xylene for 30 min each. The dehydrated samples were immersed in liquid paraffin at 65°C for 60 min, embedded, and cured overnight. Samples were cut into 5 µm-thick sections using a CUT5062 microtome (Slee, Mainz, Germany). The sections were then dried overnight at 37°C to remove any excess water. Next, sections were rehydrated through the same ethanol series for 5 min each and stained using alcian blue and periodic acid/Schiff’s reagent (Morphisto, Offenbach am Main, Germany). Finally, the sections were dehydrated again through the ethanol series, cleared in xylene, and mounted on glass slides with a coverslip. The slides were then observed under a light microscope (LSM900, Carl Zeiss Microscopy, Jena, Germany).

### Layer thickness measurement

The thickness of cell layers of the different models was assessed by acquiring images of histological sections using a 20x objective. Images were imported into FIJI^84^ and measurements were performed on three independent cultures with 10 acquired images and 10 measurements per image, resulting in n = 300 measurements per sample.

### Scanning electron microscopy

To obtain more detailed information about surface topography, scanning electron microscopy (SEM) was performed using a ZEISS EVO 10 scanning electron microscope (Carl Zeiss Microscopy, Jena, Germany) with an acceleration voltage of 6 kV. Before imaging, all samples were fixed and dehydrated in an ascending ethanol series and subsequently dried using an EM CPD300 critical point dryer (Leica, Wetzlar, Germany). Samples were sputter-coated with a thin layer of palladium/gold using an SC7620 Mini Sputter Coater (Quorum Technologies, Lewes, UK) for six minutes.

### Topography measurement

Surface morphology of the *in vitro* tissues was visualized using profilometry based on white light reflectance. Samples for SEM analysis were analysed using the topography feature of an Alpha300R^+^ confocal Raman microscopy system (WITec, Ulm, Germany). Acquired z-values representing the surface topography were exported as txt files and subsequently plotted using a custom script in MatLab (version R2023a, MathWorks, Natick, USA).

### Mucus quantification

For the quantification of mucus in *in vitro* models, cells were seeded in 12-well plates (Greiner Bio-One, Kremsmünster, Austria) and cultured as described above. For *in situ* alcian blue staining, the supernatant was carefully removed, and cell layers were fixed with methanol and acetone (1:1) for 15 min at −20°C. Samples were incubated for further 15 min with 1% alcian blue pH 2.5 (Morphisto, Offenbach am Main, Germany) and afterwards washed with ultrapure water (Veolia Water Technologies, Celle, Germany) to remove unbound dye. After complete drying of the samples, absorbance was measured at 630 nm in 225 different positions within one sample using a plate reader (Tecan, Männedorf, Switzerland). For each data point, 3 biological replicates (n=3) in 3 independent experiments (N=3) were measured. The amount of mucus is given as the relative amount after normalisation to the statically cultured Caco-2 monoculture in each independent experiment. Photographs of exemplary cultures were acquired using an EOS M50 camera (Canon, Tokio, Japan).

### Transepithelial electrical resistance (TEER) measurement

To monitor the barrier formation of *in vitro* models over the course of the 21-day cultivation period, the TEER was measured at days 7, 14, and 21 using a Millicell volt-ohm-meter (Merck, Darmstadt, Germany). Averaged blank values of empty cell culture inserts were subtracted from resistance measurements and normalised to the area of the cultivation area (0.9 cm^2^). Six independent experiments (N=6) with at least three biological replicates (n=3) were conducted.

### Immunofluorescent staining

To visualise the formation of tight junctions, *in vitro* cultures were washed twice with PBS and fixed using equal parts of ice-cold acetone and methanol for 20 min at −20°C. Samples were rinsed afterwards with PBS, blocked with 3% bovine serum albumin/1% goat serum in PBS, and incubated with an anti-zonula occludens 1 (ZO-1) antibody (1:400, #61-7300, Invitrogen, Waltham, US). After washing, samples were incubated with a goat-anti-rabbit Alexa Fluor™ 488 antibody (1:500, #A-11008, Invitrogen, Waltham, US) for 1 h at room temperature, followed by staining with Alexa Fluor™ 546 Phalloidin for 1 h (1:400, #A22283, Invitrogen, Waltham, US) and 4’,6-Diamidin-2-phenylindol, Dihydrochlorid (DAPI) for 5 min (1:100, #D1306, Invitrogen, Waltham, US). Samples were mounted on glass slides and z-stacks were acquired using an LSM900 confocal laser scanning microscope (Carl Zeiss, Jena, Germany). Z-stacks were displayed using the “surface” function of the ZEN blue edition software (Carl Zeiss, Jena, Germany). ZO-1 staining was performed for 1 sample (n=1) in 3 independent experiments (N=3), of which only exemplary images were provided in this study.

### Permeability assessment

To further assess the barrier integrity of *in vitro* models, permeability studies were conducted by measuring the flux of fluorescein isothiocyanate (FITC) labelled 4 kDa dextran (TdB Labs, Uppsala, Sweden) across the epithelia. Cell culture models were pre-incubated for 30 min with Hank’s balanced salt solution (HBSS) before TEER measurements. The basolateral compartment was replaced with fresh HBSS, and a solution of 250 µg/mL FITC-dextran was applied to the cell layer in the apical compartment. 100 µL samples were collected at time points 1, 2, 3, and 4 h. 100 µL of fresh HBSS was added to each sample’s basolateral compartment after a sample was collected. Fluorescence intensity was immediately measured in black 96-well plates (Greiner Bio-One, Kremsmünster, Austria) using a plate reader (Tecan, Männedorf, Switzerland). After the collection of the last samples, TEER values were measured again to validate the barrier stability during the course of the experiment. The apparent permeability coefficients (P_app_) were calculated using equation (1):

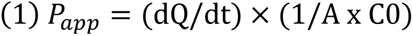

Where dQ/dt is the rate of cumulative drug transport (μg/s), A is the surface area of the cell monolayer (cm²), and C0 is the initial concentration of the drug in the donor compartment (μg/mL). Permeability tests were conducted in 3 biological replicates (n=3) in 3 independent experiments for 4 kDa FITC-dextran (N=3).

### Nanocarrier preparation

PLGA nanocarriers were prepared using the double emulsion solvent evaporation method. Initially, a solution of Resomer® RG 502H (herein referred to as PLGA) (Evonik, Darmstadt, Germany) in chloroform was combined with tris(hydroxymethyl)aminomethane buffer (tris buffer) (Sigma Aldrich, Darmstadt, Germany) and subjected to sonication (Vibra Cell™, VCX750, Sonics & Materials, Inc., USA) to generate a primary emulsion. Subsequently, the primary emulsion was added to a solution of Tween® 80 (Sigma Aldrich, Darmstadt, Germany) in ultrapure water, which was sonicated (40 %, 120 s) to produce the secondary emulsion. Solid nanocarriers were formed by evaporating the chloroform at room temperature under reduced pressure. To encapsulate Coumarin 6 (Sigma Aldrich, Darmstadt, Germany), the payload was dissolved in the chloroform phase before nanocarrier production. The resulting nanocarriers were then subjected to centrifugation at 21500 xg for 20 minutes to separate any non-encapsulated drug molecules. The supernatant was discarded, and the NCs were subjected to sonication and redispersion in ultrapure water. This washing process was repeated three times.

PEG-PLGA nanocarriers were produced accordingly, with a 3:1 mixture of PLGA and PEG-PLGA (Sigma Aldrich, Darmstadt, Germany) as polymers. Finally, the nanocarriers were stored at 8 °C in the absence of light until further use.

Hydrodynamic diameters and PDI values of the formulations were determined using dynamic light scattering, while the zeta-potential was measured using electrophoretic light scattering (Zetasizer NanoZS, Malvern Panalytical, Malvern, UK).

Payload leakage was assessed in HBSS at 37 °C. Samples were drawn after 0.5, 1, 3, 5, 8, and 24 h incubation and centrifuged (21500 xg, 4 °C, 20 min) (Thermo Scientific SL 8R Centrifuge, Thermo Fisher Scientific, Darmstadt, Germany). The fluorescence intensity of the supernatant was measured measured (485 nm excitation, 535 nm emission) (Spark multimode microplate reader, Tecan, Männerdorf, Switzerland) to determine the concentration of leaked coumarin 6.

### Cytotoxicity testing

The cytotoxic potential of nanocarrier formulations was evaluated in the different *in vitro* models following a 21-day cultivation under static or dynamic conditions in 48 well plates. Nanocarriers were applied at a final concentration of 0.2 mg/mL in HBSS. As a positive control for each tissue model, pure HBSS without nanocarriers was applied. *In vitro* models were incubated for 6 h, followed by the addition of 1 mg/mL MTT reagent (3-(4,5-Dimethylthiazol-2-yl)-2,5-diphenyltetrazolium bromide) (Sigma Aldrich, Darmstadt, Germany). After the incubation period of 4 h, the culture medium was removed, and the resulting formazan crystals were dissolved in dimethyl sulfoxide (DMSO). Absorbance measurements were performed at a wavelength of 570 nm using a microplate reader (Tecan, Männedorf, Switzerland), allowing for the quantification of cell viability and the assessment of nanocarrier-induced cytotoxicity via equation (2)^85^:

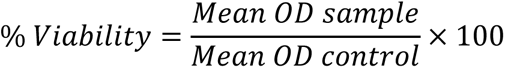

### Nanocarrier permeation testing

The permeation of PLGA and PEG-PLGA nanocarriers was assessed analogously to permeability studies described above. Nanocarriers were applied at a concentration of 0.2 mg/mL and samples were collected at time points 1, 2, 3, 4, and 6 h. The permeation of nanocarrier formulations was assessed in 3 biological replicates (n=3) in 3 independent experiments (N=3). For each independent experiment, 1 sample was fixed with a 3.7% buffered paraformaldehyde solution and stained using Alexa Fluor™ 546 Phalloidin and (DAPI) as described above.

### Statistical analysis

Statistical analysis was performed using R: A Language and Environment for Statistical Computing (R Foundation for Statistical Computing, Vienna, Austria) and applied for evaluation of the TEER value, FITC-dextran leakage, nanocarrier permeation assays, mucus quantification, and layer thickness. After testing of normality distribution (Shapiro-Wilk test) and variance homogeneity (Bartlett test), the presence of statistically significant differences (alpha level 0.05) between groups was determined either by one-way analysis of the means followed by Welch test with Bonferroni correction or by ANOVA and Tukey multiple comparisons of means (95% family-wise confidence level). P values < 0.05 were considered statistically significant (* p < 0.05, ** 0.05 < p < 0.01, *** 0.01 < p < 0.001).

## Supporting information

Supplemental information (all figures)

## Author information

### Authors

Jonas Schreiner – Institute of Pharmaceutical Technology, Goethe University Frankfurt, Frankfurt am Main, Germany;

Florentin Baur – Institute of Pharmaceutical Technology, Goethe University Frankfurt, Frankfurt am Main, Germany;

Sarah Vogel-Kindgen – Institute of Pharmaceutical Technology, Goethe University Frankfurt, Frankfurt am Main, Germany;

## Acknowledgement

This work was conducted in the framework of the EUbOPEN project, which has received funding from the Innovative Medicines Initiative 2 Joint Undertaking under grant agreement No 875510. This study was supported by the Cluster project ENABLE funded by the Hessian Ministry for Science and the Arts. This work was supported by PROXIDRUGS as part of the initiative “Clusters4Future”, funded by the Federal Ministry of Education and Research BMBF (03ZU1109XX). Schemes were, in part, created with BioRender.com.

