## Supplemental information (all figures) for "Predicting nanocarrier permeation across the human intestine *in vitro*: Model matters"

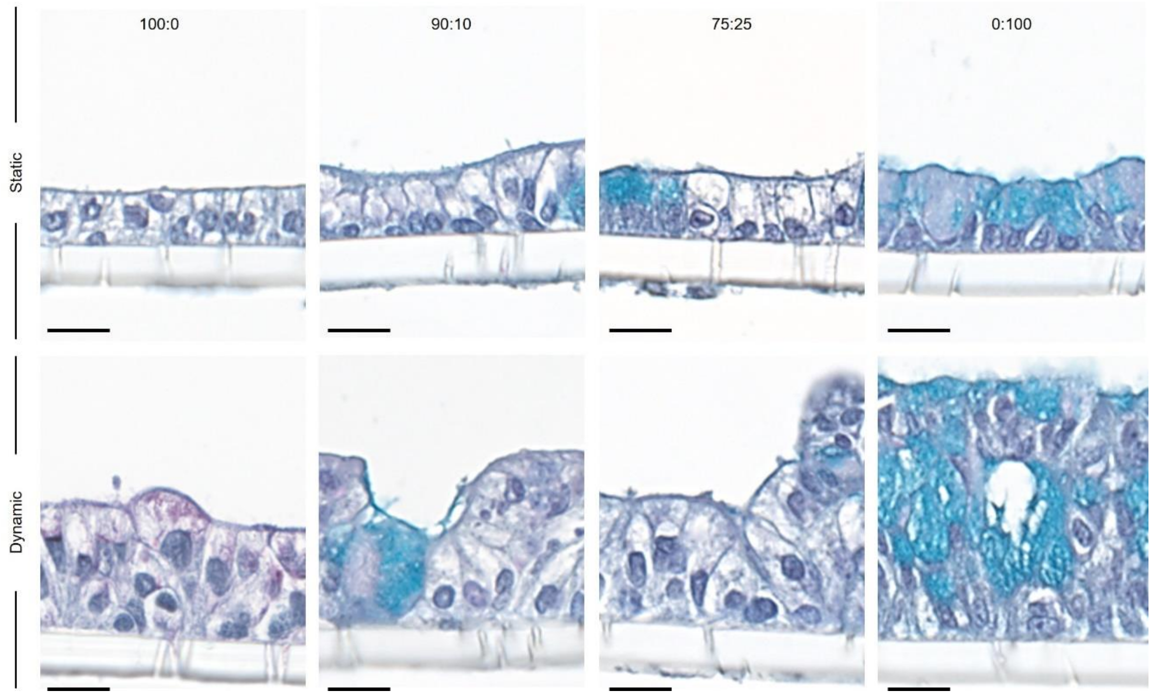


**Figure SI1**. Magnification of alcian blue/PAS staining of histological sections of *in vitro* tissue models cultured under static or dynamic conditions. Scale bars: 20 µm.

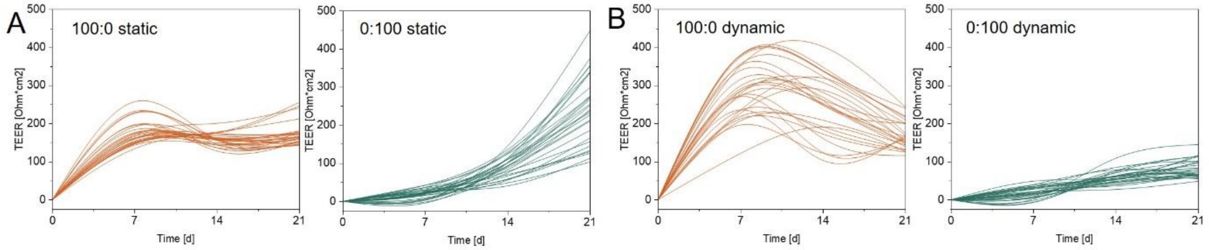


**Figure SI2**: Development of transepithelial electrical resistance (TEER) of Caco-2 (100:0) and HT29-MTX (0:100) monocultures over 21 days. Each line in the spline graphs represents a single culture of the respective *in vitro* model.

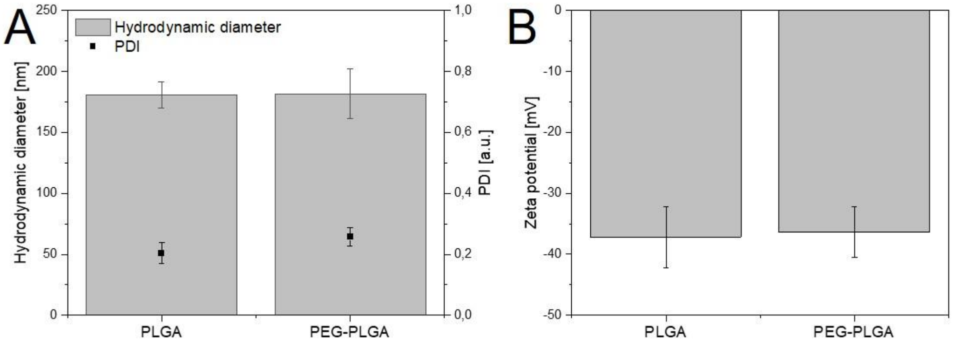


**Figure SI3**. Nanocarrier hydrodynamic diameter (A) and surface charge (B).

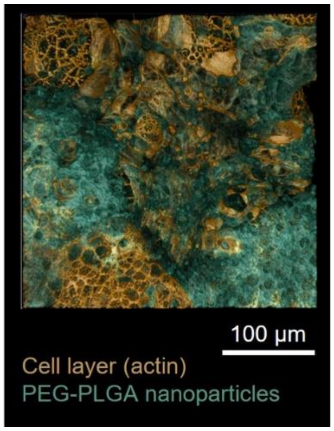


**Figure SI4**. Confocal fluorescence image of dynamic 75:25 culture after transport study with PEG-PLGA nanocarriers.

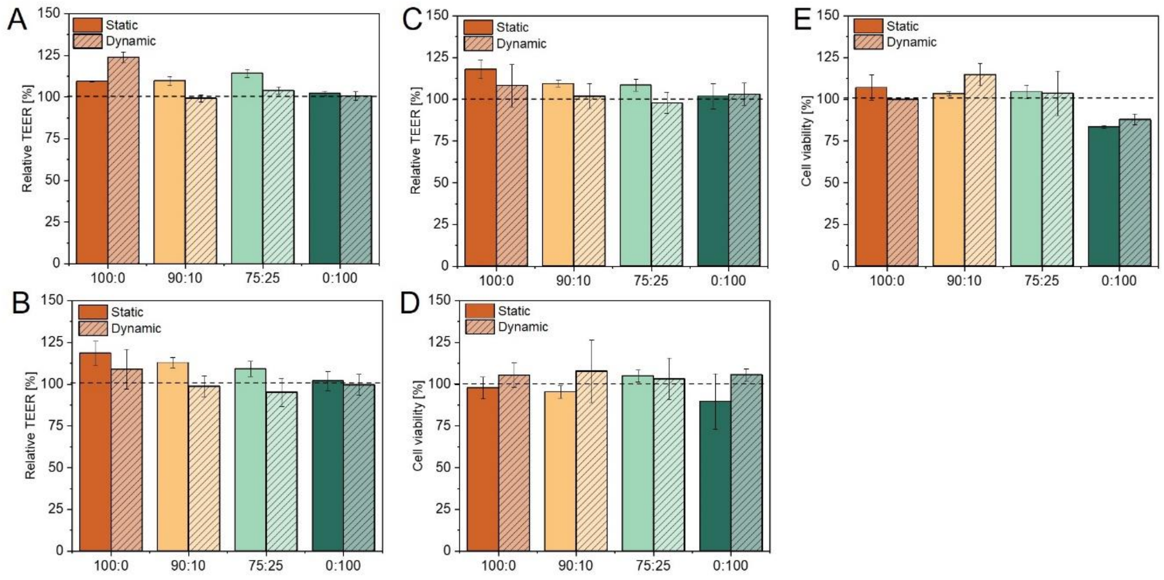


**Figure SI5**. Barrier integrity as a function of relative transepithelial electrical resistance (TEER) after permeation testing of 4 kDa FITC-dextran (A), PLGA nanocarriers (B), and PEG-PLGA nanocarriers (C). Cell viability of different *in vitro* models after 24 h incubation with PLGA nanocarriers (D) and PEG-PLGA nanocarriers (E).

### Author information

#### Corresponding Authors

Nathalie Jung – Institute of Pharmaceutical Technology, Goethe University Frankfurt, Frankfurt am Main, Germany;

Maike Windbergs – Institute of Pharmaceutical Technology, Goethe University Frankfurt, Frankfurt am Main, Germany;

#### Authors

Jonas Schreiner – Institute of Pharmaceutical Technology, Goethe University Frankfurt, Frankfurt am Main, Germany;

Florentin Baur – Institute of Pharmaceutical Technology, Goethe University Frankfurt, Frankfurt am Main, Germany;

Sarah Vogel-Kindgen – Institute of Pharmaceutical Technology, Goethe University Frankfurt, Frankfurt am Main, Germany;
